# Infection of mouse neural progenitor cells by *Toxoplasma gondii* affects *in vitro* proliferation, differentiation and migration

**DOI:** 10.1101/2021.05.31.446482

**Authors:** Luiza Bendia Pires, Helene Santos Barbosa, Marcelo Felippe Santiago, Daniel Adesse

**Author notes:** **Corresponding Author:** Daniel Adesse, Laboratório de Biologia Estrutural, Avenida Brasil, 4365, Pavilhão Carlos Chagas, sl. 307, Rio de Janeiro, RJ, Brazil, **Postal code:** 21040-360, **E-mail:**, **Alternative e-mail:**.

## Abstract

Congenital toxoplasmosis constitutes a major cause of pre- and post-natal complications. Fetal infection with *Toxoplasma gondii* influences development and can lead to microcephaly, encephalitis, and neurological abnormalities. Few studies have attempted to explain the impact of *T. gondii* infection on the physiology of mature nerve cells, and no systematic study concerning the effect of infection of neural progenitor cells by *T. gondii* in the biology of these progenitors is available. We infected cortical intermediate progenitor cell cultivated as neurospheres obtained from E16.5 Swiss Webster mice with *T. gondii* (Me49 strain) tachyzoites to mimic the developing mouse cerebral cortex *in vitro*. Infection decreased cell proliferation as detected by Ki67 staining at 48 and 72 hours post infection (hpi) in floating neurospheres, resulting in reduced cellularity at 96 hpi. Neurogenic and gliogenic potential, assessed in plated neurospheres, was shown to be impaired in infected cultures, as indicated by neurofilament heavy chain (NF-200) and GFAP staining, respectively. To further investigate the impact of infection on neuronal differentiation, Neuro2a neuroblasts were infected and after 24 hpi, neurogenic differentiation was induced with serum withdrawal. We confirmed that infection induces a decrease in neuroblast-neuron differentiation rates in cells stained for NF-200, with reduced neuritogenesis. Migration rates were analyzed in plated neurospheres. At 120 h after plating, infected cultures exhibited decreased overall migration rates and altered the radial migration of nestin-, GFAP- and NF-200-positive cells. These findings indicate that *T. gondii* infection of neural progenitor cells may lead to reduced neuro/gliogenesis due to an imbalance in cell proliferation alongside an altered migratory profile. If translated to the *in vivo* situation, these data could explain, in part, the cortical malformations observed in congenitally infected individuals.

## INTRODUCTION

Toxoplasmosis is one of the most common zoonotic diseases worldwide. It is estimated that 1/3 of the world’s human population is latently infected by *Toxoplasma gondii (T. gondii)*. In healthy individuals, primary infection by *T. gondii* causes mild symptoms, whereas in the immunocompromised patients or in the developing fetus, this parasite can cause life-threatening infections with severe neurological and ocular manifestations (Desmonts and Couvreur, 1974; Luft et al., 1993; Roberts and McLeod, 1999).

In most hosts, *T. gondii* establishes a life-long, latent infection in certain tissues, such as skeletal and cardiac muscles, or the central nervous system (CNS), which includes the brain, the spinal cord, and the retina (Wohlfert et al., 2017). *T. gondii* transmission occurs mainly via the oral route, through the ingestion of cysts containing bradyzoites or oocysts containing sporozoites or by congenital transmission (Tenter et al., 2000; Montoya & Liesenfeld, 2004). The parasite rapidly infects host cells, differentiating to the fast replicating tachyzoite form and dividing intracellularly by a process termed endodyogeny (Sheffield and Melton, 1968; Dubey et al. 1998), followed by host cell egress, promoting cell lysis (Blader et al. 2015). During the initial weeks of infection, tachyzoites can be found in the brain parenchyma and in migrating dendritic cells (Harker et al., 2015; Cabral et al., 2016; Estato et al., 2018). As infection proceeds, *T. gondii* transitions into the chronic stage of infection via conversion to the slowly replicating bradyzoite, which encysts. Several transitions occur between tachyzoites and bradyzoites/cysts, some of which are thought to enable the bradyzoite/cyst to escape immune detection, thereby leading to persistent infection (Kim & Boothroyd, 2005). In humans and rodents, the brain is the major encystment and persistence organ (Remington & Cavanaugh, 1965).

Vertical transmission can occur when a seronegative pregnant female is infected (Hall, 1992). The parasite can reach the fetus and infect the brain, where both tachyzoites and bradyzoites can be found (Ferguson et al., 2013).

Congenital infection by *T. gondii* can lead to aggressive manifestations, including blindness, retinochoroiditis, cerebral calcifications, hydrocephalous, ventriculomegaly and microcephaly (see Campos et al (2020) for reviews). Experimentally, *T. gondii* is able to infect and form cysts in neurons, astrocytes and microglial cells (Lüder et al., 1999). *In vivo* reports, however, have shown that the parasite preferentially infects neuronal cells where tissue cysts are formed and, to a lesser extent, astrocytes (Melzer et al., 2010; Cabral et al., 2016).

The incidence and severity of congenital toxoplasmosis infection depends on the gestational period when infection occurs. The risk of vertical transmission increases over the gestational weeks, of 15% in the 13^th^ week, 44% in the 26^th^ and 71% in the 36^th^, increasing to 90% in the last week of pregnancy. However, the severity of fetal damage is inversely proportional to the infection period. Severe manifestations are seen in the offspring of women who acquired the infection during the beginning of the pregnancy, whereas it may be subclinical in neonates born to mothers infected at the end of the pregnancy (Hall, 1992). Therefore, *T. gondii* is included in the TORCH complex, comprising pathogens that, when acquired during pregnancy, can migrate to the fetus and cause neurological malformations, including microcephaly (see Campos et al, 2020, for reviews).

Microcephaly is a condition in which patients exhibit a marked decrease in the size of the head and the brain (>2 standard deviations of the mean head size for the same age and gender). Although cortical organization is mostly preserved in the smaller brain, patients may display significant intellectual disability (Jayaraman et al., 2018; Devakumar et al., 2018). Microcephaly is usually associated with a decrease in progenitor cell numbers, which may be due to decreased proliferation, changes in symmetric and asymmetric division patterns and increased progenitor cell death (Barkovich et al., 2005; Passemard et al., 2013). In the case of acquired congenital microcephalies caused by gestational exposure to teratogenic agents, brain damage can lead to macroscopic (malformations, disruption) or microscopic (dysplasia) CNS developmental anomalies (Passemard et al., 2013, for reviews).

Since microcephaly is observed in congenital toxoplasmosis, we aimed at developing an *in vitro* system comprising neural progenitor cell (NPC) infection to study the outcomes of *T. gondii* infection on cell proliferation, multipotency and migration. Using a primary culture of NPCs as floating neurospheres, we were able to determine that *T. gondii* reduces NPC proliferation rates, which leads to reduced cellularity. When plated onto a substrate and cultivated with specific culture medium, infected cultures exhibited reduced astrocytogenesis and neurogenesis, as well as impaired migration potential.

## METHODS

### 1. Cell culture

Mouse neural progenitor cells were isolated following protocols described by Duval et al., (2002) and Santiago et al., (2010). The anterior encephalons of 10-13 E16.5 Swiss Webster mouse embryos were aspirated with a 22G needle into a syringe containing sterile PBS. The tissue was homogenized with a Pasteur pipette in Growth medium [3 ml DMEM-F12 with 2% of B27 without vitamin A (Gibco), 20 ng.mL^-1^ Epidermal Growth Factor (EGF, Sigma Aldrich), 16 mM glucose, GlutaMax (Thermo Fisher), 4.8 mM glucose and an antibiotic solution (Thermo Fisher)]. Cells were centrifuged at 300 rpm for 7 minutes to remove dead cells and then centrifuged again at 1,200 rpm for 7 minutes, resuspended in 5 mL of growth medium and plated on T25 culture flasks. The cultures were maintained at 37°C under a 5% CO_2_ atmosphere for seven days, with the addition of 1 µL of EGF at 20 ng.mL^-1^ every two days. The neurospheres were centrifuged and dissociated in a 0.25% trypsin and 0.01% EDTA solution and viability was assessed by Trypan blue stain exclusion. A total of 2.8 x 10^5^ cells were plated in T25 flasks and cultivated for fourteen days until another round of dissociation. For the experiments, 5 x 10^4^ second passage NPCs were plated in 5 mL of the growth medium in T25 flasks. After four days in culture, the cells were infected with the tachyzoite forms of *T. gondii*.

Neuro2a cells were a kind gift from Prof. Fiona Francis and Prof. Richard Belvindrah (Institut du Fer a Moulin, INSERM Paris) and maintained in a Proliferation Medium [**PM**, DMEM-F12 medium supplemented with 10% fetal bovine serum (FBS) and 1% of an antibiotic solution (Streptomycin/Penicillin, ThermoFisher)]. The medium was replaced every 48-72 hours. For differentiation induction, the PM was replaced with Differentiation Medium [DM, DMEM-F12 (Gibco) supplemented with 0.1% bovine serum albumin (BSA, Sigma Aldrich) and 1% of an antibiotic solution (Thermo Fisher)] and maintained at 37°C under a 5% CO_2_ atmosphere.

### 2. Parasites

Parasites of the Me49 strain were obtained from brains of C57BL/6 mice infected 45 days before isolation. Cysts were ruptured with an acid pepsin solution and free parasites were added to Vero cell monolayers (ATCC). After two weeks of culture re-infections, tachyzoites released from the supernatant were collected and centrifuged prior to use. For the experiments, cultures were infected with tachyzoites at a MOI of 1 for two hours. One T25 with NPCs was dissociated and counted at each experiment in order to estimate the number of host cells per flask. After 24 hours of infection, cells were washed in Ringer’s solution and fresh growth medium was added.

### 3. Experimental design

For the neurospheres assays, two approaches were applied: (1) floating neurospheres were infected after two days of plating in a MOI of 1. The MOI was calculated based on the number of neurospheres per well, by preparing an extra well used to dissociate and count the number of viable cells. After 24-96 hours of infection, cultures were fixed and processed for whole mount immunofluorescence; (2) floating neurospheres were infected after two days of plating using a MOI of 1 and, after 24 hours of infection, uninfected and infected cultures were plated separately in a final volume of 200 µL of NPC differentiation medium [DMEM-F12, supplemented with 1% B27 with retinoic acid (Gibco, Life Technologies),1% GlutaMAX™ (Gibco, Life Technologies), 2% N2 supplement (Gibco, Life Technologies) e 1% antibiotic solution Penicillin/Streptomycin (Gibco)] onto glass coverslips previously coated with fibronectin and poly-L-lysine (Sigma Aldrich). After two hours of plating, an additional 300 µL of NPC differentiation medium was added to the wells and cultures were photographed under an inverted light microscope with phase contrast illumination (considered T = 0 h). After 24, 48 and 120 hours of plating, cultures were photographed and fixed at 48 and 120 h for immunofluorescence assays.

Neuro2a cells were plated onto glass coverslips previously coated with poly-L-lysine in borate buffer, washed extensively with tri-distilled water prior to plating and maintained in PM. After 24 hours of plating, cells were infected with *T. gondii* tachyzoites (Me49 strain) at a multiplicity of infection of 3 parasites per cell (MOI 3). At 24 hours after infection, the medium was changed to DM in half of the cultures and were fixed after 24 hours of differentiation induction, corresponding to 48 hours of infection (48 hpi).

### 4. Whole-mount immunofluorescence

Neurospheres were collected, washed in PBS and fixed in 4% paraformaldehyde (Sigma Aldrich) for 5 minutes at 20 °C, permeabilized with a 0.5% Triton x-100 (Sigma Aldrich) solution in PBS, blocked with a 4% BSA solution for 30 minutes and incubated with primary antibodies against SAG1 (tachyzoite marker, mouse antibody, Santa Cruz Biotechnology) and Ki67 (rabbit polyclonal antibody, ABCAM) overnight at 4 °C. Secondary donkey anti-rabbit IgG AlexaFluor 488 (Thermo Fisher) and goat anti-mouse IgG AlexaFluor 594 (Thermo Fisher) antibodies were incubated for one hour at 37 °C. Nuclei were visualized by incubating cells with DAPI (4′,6-Diamidine-2′-phenylindole dihydrochloride, Sigma Aldrich) at 0.2 mg/mL for 5 minutes at 20 °C. Finally, the neurospheres were transferred to a DABCO (1,4 Diazabicyclo[2.2.2]octane, Sigma Aldrich) solution containing 50% glycerol in PBS and visualized under a Zeiss Meta 510 confocal microscope.

### 5. Immunostaining in attached cells

Neuro2a or NPCs were fixed at desired times in 4% paraformaldehyde for 20 minutes at 4 °C. Cells were then permeabilized with Triton x-100 0.5% for 30 minutes, blocked with BSA 3% and incubated with primary antibodies overnight at 4 °C. The antibodies used herein are listed in **Table 1**. Cells were washed with PBS and incubated in secondary antibodies coupled to AlexaFluor for 1 h at 37 °C. Nuclei were visualized by incubating cells with DAPI at 0.2 mg/mL for 5 minutes at 20 °C. Slides were mounted in a DABCO solution and visualized under a Zeiss Meta 510 confocal microscope.

**Table 1:** Primary and secondary antibodies used for immunocytochemistry assays.

| Antibody | Producer | Source | Dilution |
| --- | --- | --- | --- |
| Anti-Ki67 | ABCAM | Rabbit | 1:80 |
| **Anti-SAG1** | Santa Cruz | Mouse | 1:100 |
| **Anti-β-III-tubulin** | Cell Signaling Technologies | Mouse | 1:200 |
| **Anti-nestin** | Millipore | Mouse | 1:200 |
| **Anti-GFAP** | DAKO | Rabbit | 1:500 |
| **Anti-neurofilament-200** | Sigma | Rabbit | 1:300 |
| **Anti-NeuN** | Millipore | Mouse | 1:300 |
| **Anti-mouse IgG (Alexa Fluor 594)** | Life Technologies | Goat | 1:200 |
| **Anti-rabbit IgG (Alexa Fluor 488)** | Life Technologies | Goat | 1:2,000 |

### 6. Scanning Electron Microscopy

Floating neurosphere were washed twice in PBS and fixed in 2.5% glutaraldehyde (Sigma Aldrich) for 1 hour at 4 °C, followed by washes in cacodylate buffer. Post-fixation was performed with osmium tetroxide (Sigma Aldrich) 0.1 M diluted in cacodylate buffer for 1 hour at 4 °C and samples were then dehydrated in an ascending ethanol series. Samples were placed on top of round glass coverslips coated with adhesive tape and mounted on aluminum stubs, dried by the critical point method, coated with a ∼20-nm layer of gold by ion sputter coater (Cressington Sputter Coater 108) and analyzed using a Jeol JSM 6390LV scanning electron microscope at 15 kV, at the Rudolf Barth Electron Microscopy Facility, Oswaldo Cruz Institute (IOC), Fiocruz.

### 7. Morphometric analyses

Neurospheres were photographed using an inverted Zeiss Axiovert microscope at 10 or 20x magnification. Surfaces were determined by calculating the radius of each neurosphere and applying the formula A=4πR^2^, where A is the area and R is the radius of the neurosphere. At least ten neurospheres from each culture at each time point from two independent experiments were counted. To determine the migrated area in attached neurospheres, at least 25 neurospheres in each experimental condition from at least three primary cell preparations were measured using the ImageJ software. To determine neurite growth measurements, β-III-tubulin- and neurofilament-stained Neuro2a cells were imaged using a confocal microscope and maximal projection of the whole Z axis was obtained with the Zen software (Zeiss). Neurite lengths were determined by the fiji plugin (ImageJ). The number of nestin-, GFAP- and neurofilament-positive filaments in migrated neurospheres was determined by tracing a virtual line in each neurosphere, placed on the margin where the migratory halo was established and counting the number of filaments that crossed that line. The number of positive filaments of each neurosphere was normalized by the length of the virtual line. Total numbers of β-III-tubulin-positive cells were determined by counting the number of stained cells in the whole neurosphere and dividing this value by the neurosphere-occupied area. A Sholl analysis was performed in nestin-, GFAP- and NF-200-labeled cultures 48 and 120 h after plating, by counting the number of intersects of the cell filaments with spheres of varying radii centered in the neurosphere’s core. The number of intersects of positively-labeled cells was plotted against the sphere-radius. The Sholl analysis allowed for the derivation of a number of parameters that further describe the complexity of migrated neurospheres for every culture, as follows: the mean value (MV) is the average number of intersects over all radii, the critical value (CV) is the maximum number of intersects and the critical radius (CR) is the radius at which the CV occurs. The maximum distance (MaxDist) is the radius at which no more intersects occur (Malinowski et al., 2019).

### 8. Statistical analyses

Data were obtained from three independent experiments in Neuro2a cells and at least three independent neurosphere experiments. At least 25 floating neurospheres per experimental condition were analyzed for Ki67-based proliferation analysis and a minimum of 15 adhered neurospheres per experimental condition were used for the migration area analysis. The results were analyzed using the GraphPad Prism software version 8.4.3, through unpaired Student’s T test or Two-Way ANOVA with Bonferroni post-test. Results were considered statistically significant when p<0.05.

### 9. Animal work

Swiss Webster and C57BL/6 mice were obtained from the Instituto de Ciência e Tecnologia em Biomodelos (Fiocruz). Pregnant SW female mice at E16.5 were euthanized immediately for the NPC preparations. C57BL/6 mice were kept at the Animal Facility of the Instituto Oswaldo Cruz with food and water ad libitum and kept at 20 °C. At least two days after arrival, five animals received 50 cysts of *T. gondii*, ME49 strain, via IP route in 100 µL of PBS. After 45 days of infection, animals were euthanized and brains were surgically removed and chopped in PBS solution using sterile scissors. Brain suspensions were passed through different needles using sterile syringe with caliber up to 26G. Cysts were counted in 20 µL of suspension and kept at 4 °C for up to a month. To obtain free bradyzoites, cysts were ruptured with acid pepsin solution and then added to uninfected cultures of Vero cells. Use of animals was approved by the Commission of Ethics in the Use of Laboratory Mice (CEUA-IOC) from the Oswaldo Cruz Institute under the license number L-048/2015.

## RESULTS

### 1. *T. gondii* infection impairs neural progenitor cell proliferation in floating neurospheres

Floating neurospheres were infected with *T. gondii* tachyzoites and analyzed 24-96 hours post infection (hpi, **Figure 1**). Scanning Electron Microscopy was used to visualize general morphological neurosphere aspects (**Figure 1B**). At 96 hpi, infected cultures exhibited increased cell processes on their surface (right panels), when compared to uninfected cultures (left panels) cultured for the same time (**Figure 1B**). To detect proliferating cells, whole neurospheres were immunostained for Ki67 and counterstained with DAPI (**Figure 1C-D**). At 24 hpi, corresponding to 48 hours of plating, uninfected neurospheres exhibited 45±21% of Ki67-positive cells, and infected neurospheres, 47±18% (p>0.05). However, *T. gondii*-infected displayed 14% and 11% reductions (p<0.01, two-way ANOVA with Bonferroni post-test) in Ki67-positive cells at 48 and 72 hpi, respectively when compared to their time-matched controls (**Figure 1C** and **D**). Intracellular tachyzoites were detected with anti-SAG1 antibody, as shown in red in **Figure 1C**, and were more noticeable in infected cultures after 48 hpi. In order to determine whether reduced proliferation would be reflected in the overall cellularity of the neurospheres, we counted the number of DAPI-positive cells in the longest diameter of each neurosphere, by confocal microscopy (**Figure 1E**). *T. gondii* infection led to a negative impact on the number of DAPI-stained nuclei at 96 hpi, since control cultures contained of 6.6 x 10^3^ ±1.9 cells/mm^2^ and *T. gondii*-infected cultures, 4.9 x 10^3^ ±1.4 cells/mm^2^ (p=0.0183, Two-way ANOVA with Bonferroni post-test), representing a 26% decrease in cellularity.

**Figure 1:**
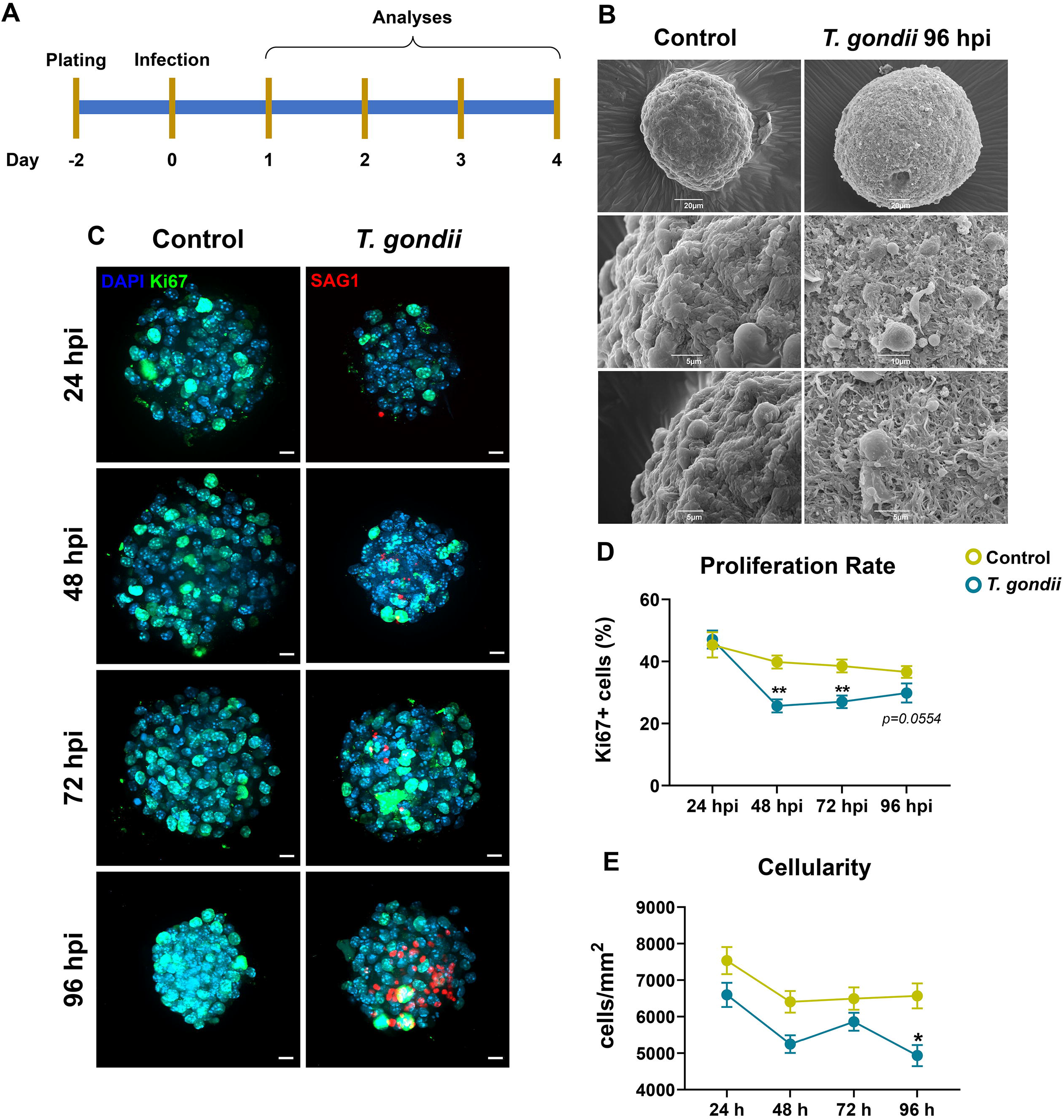
*T. gondii* affects NPC proliferation rates in floating neurospheres. Cultures were infected two days after plating with *T. gondii* tachyzoites (Me49 strain). After 24-96 hpi, cultures were assessed concerning host cell proliferation rates (experimental design in **A**). Scanning electron microscopy (**B**) was used to display the overall morphology of uninfected (left panels) and *T. gondii*-infected cultures at 96 hpi (right panels). Uninfected cultures showed a smooth surface, whereas infected dishes displayed neurospheres with a high number of cell projections. Ki67 immunostaining (green, in **C**) was used to assess proliferative cells in relation to total cell population, as determined by DAPI staining (blue). Intracellular tachyzoites were detected with SAG-1 antibody (in red) and are shown in detail in the inset. Infection led to the reduction of Ki67-positive cells at 48 and 72 hpi (**D**), and a reduction in cellularity at 96 hpi was detected by phase contrast microscopy (**E**). *: p<0.05; **: p<0.01, Two-Way ANOVA with Bonferroni post-test. Minimum of 25 neurospheres per experimental condition in **D** and **E**. Scale bars in **C** = 10 µm and 5 µm in inset.

### 2. Infected neurospheres exhibit impaired neuronal differentiation

After 24 hours of infection, floating neurospheres were plated onto glass coverslips in DM and fixed after 48 and 120 hours. In order to assess NPC neurogenic and gliogenic potential, immunocytochemistry was performed for different markers. We first quantified the number of nestin-and Ki67-positive cells, as indicators of neural progenitor cells. At 48 hours after plating, uninfected neurospheres showed an average of 64±14 nestin-positive filaments per millimeter (**Figure 2B**), similar to *T. gondii*-infected neurospheres (63±21, **Figure 2C**). At 120 hours of plating, which corresponds to 144 hours of infection, the number of nestin-positive filaments were decreased in both groups compared to what was observed at 48 h (53±14 and 58±8 filaments/mm in control and infected cultures, respectively **Figure 2D** and **E**), although no significant changes were observed between these two groups. Similarly, no significant changes were observed regarding proliferative cell rates, as indicated by Ki67 staining (**Figure 2B-E** and **G**). As expected, Ki67-positively stained cells exhibited decreased numbers at 120 h in uninfected dishes (2.1±1.7% in controls and 1.9±2.5% in infected cells) when compared to 48 h (13.1±7% and 13.7±8% in controls and infected cells, respectively), although no significant changes were observed in infected cultures compared their respective controls (**Figure 2G**). No changes were noted concerning cell death as determined by the number of pyknotic nuclei in infected cultures when compared to controls (**Supplementary Figure S1**).

**Figure 2:**
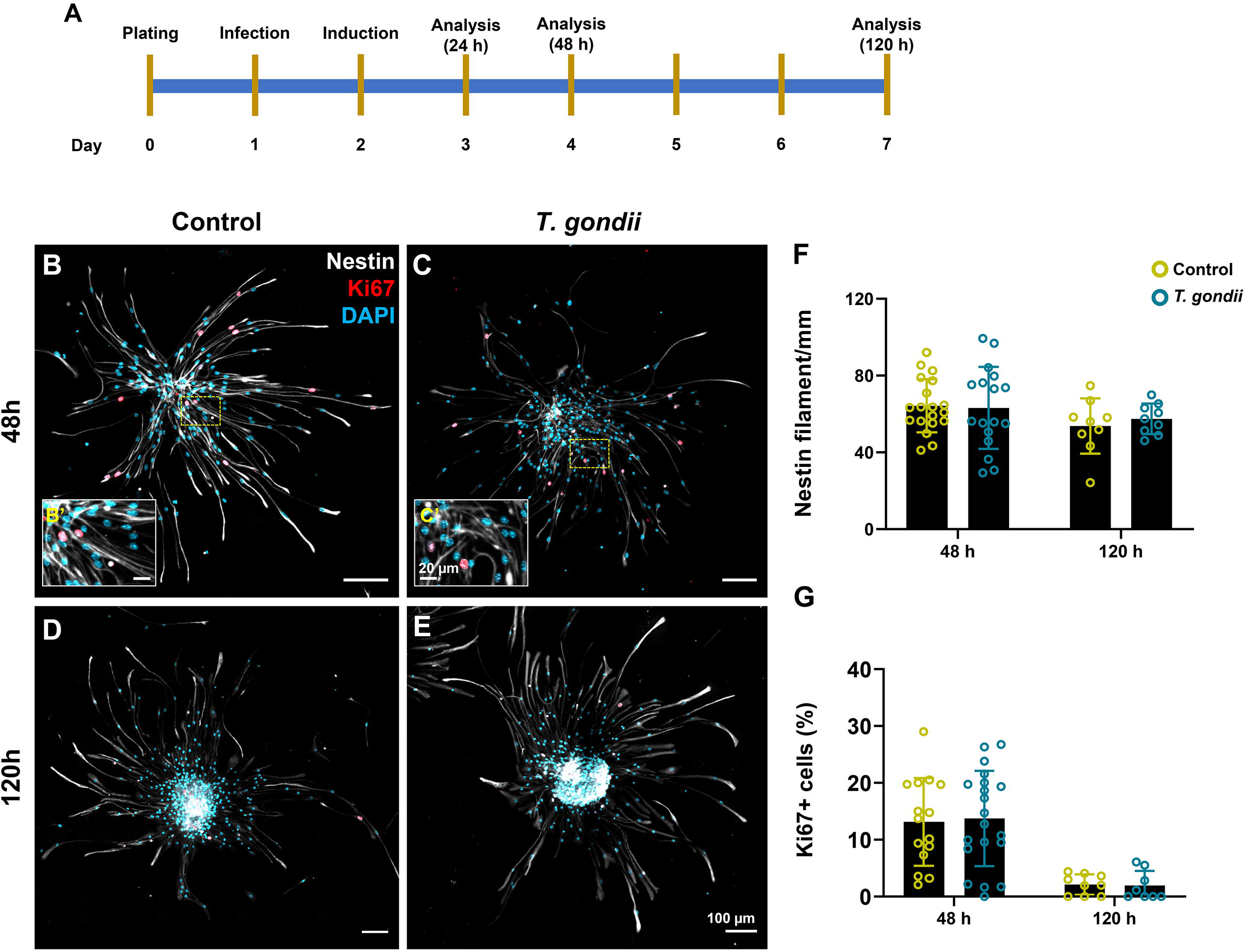
Proliferation of neural progenitor cells was not affected in infected cultures in DM. Floating neurospheres were infected in proliferation medium and, after 24 hpi, were plated onto glass coverslips in differentiation medium (the experimental design is shown in **A**). Cultures were analyzed after 48 and 120 h of plating for nestin (in white) and Ki67 (in red), neural stem cell and proliferation markers, respectively (**B-E**). Host cell nuclei were stained with DAPI (in blue). No changes were observed in nestin-positive filaments per mm (**F**) or Ki67-positive cells (**G**) in infected cultures when compared with uninfected controls. Scale bars = 100 µm in **B**-**E** and 20 µm in **B’** and **C’**.

Subsequently, we analyzed neuronal production through two different markers, β-III-tubulin and Neurofilament heavy chain (NF-200) (**Figure 3**). β-III-tubulin (TUJ1) is an early neural cell differentiation marker (Menezes & Luskin, 1994). Uninfected cultures at 48 hours of plating displayed 1.44±1.5 TUJ1-positive cells per mm^2^ whereas *T. gondii*-infected cultures contained 3.23±3.7/mm^2^ (p>0.05). After five days of plating, the overall number of TUJ1-positive cells decreased to 0.17 and 0.32 cells per mm^2^ in control and infected cultures, respectively (**Figure 3C-D** and **I**). NF-200 was used as a late neuronal differentiation marker (**Figure 3E-H**). At 48 h of plating, no changes were observed between the control and infected groups, exhibiting 30.42±20 and 43.57±22 NF-200 filaments per mm, respectively (p>0.05, **Figure 3E-F** and **J**). As expected, the number of NF-200-positive filaments was increased in uninfected cultures at 120 hours of plating (**Figure 3G-H** and **J**), reaching an average of 83.34±14 filaments/mm (p<0.0005 when compared to uninfected cultures at 48 h, Two-way ANOVA with Bonferroni post-test). This increase was impaired in *T. gondii*-infected cultures, where 54.45±25 filaments/mm of NF-200 were observed (**Figure 3J**).

**Figure 3:**
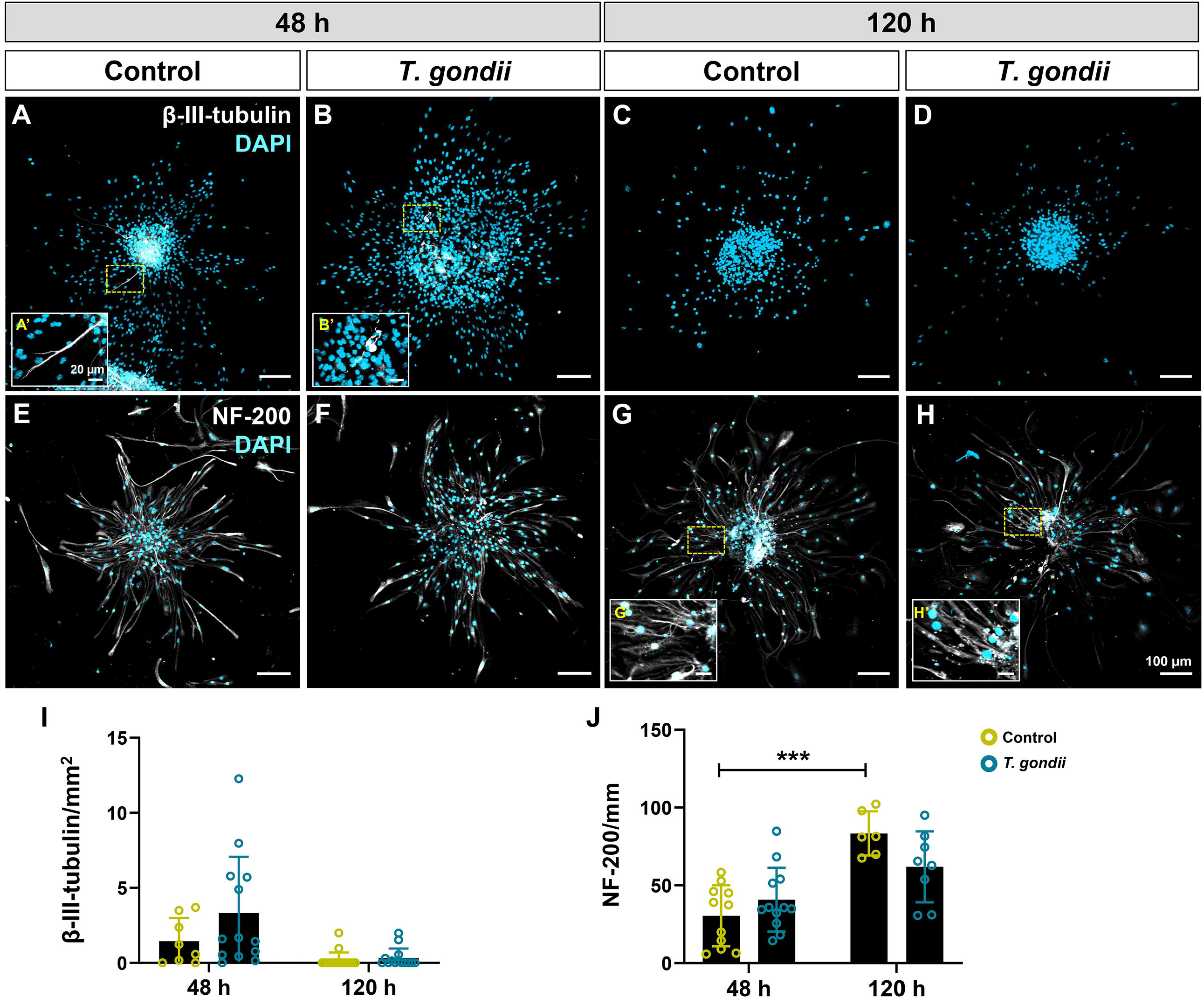
*T. gondii* infection impairs neurogenic potential in NPCs. Neurospheres were plated onto glass coverslips and analyzed after 48 and 120 hours for neuronal production by TUJ1 (**A-D**, in white) and NF-200 (**E-H**, in white) staining. Cultures were counterstained with DAPI (in blue) and analyzed by confocal microscopy. No significant changes were observed in TUJ1 stained cultures (**I**). Uninfected cultures exhibited increased numbers of NF-200 at 120 h when compared to 48 h, whereas this effect was abrogated in *T. gondii*-infected dishes (**J**). ***: p<0.001, Two-Way ANOVA with Bonferroni post-test. Scale bars = 100 µm **A**-**H** and 20 µm in **A’**, **B’**, **G’** and **H’**.

We further evaluated neuronal differentiation using Neuro2a murine neuroblasts. Cells were infected with tachyzoites in PM and, after 24 hours of infection, the medium was switched to DM in half the cultures. After 24 hours of differentiation induction, cells were stained for TUJ1 or NF-200. Neurogenesis rates were determined by the number of neuron-like cells, i.e. cells with neurites longer than the cell body stained with β-III-tubulin or NF-200 in DM divided by the number found in PM cultures. Uninfected Neuro2a cells exhibited neurogenesis rates of 1.65±0.8 and 3.8±1.4 detected by TUJ1 and NF-200, respectively (**Figure 4**). The neurogenic rate of *T. gondii*-infected Neuro2a cells was not significantly altered as revealed by TUJ1 (2.64±1.3, p=0.3, unpaired Student’s T test), but was decreased by 53% in NF-200-stained cultures (1.18±0.8, p=0.0471, unpaired Student’s T test).

**Figure 4:**
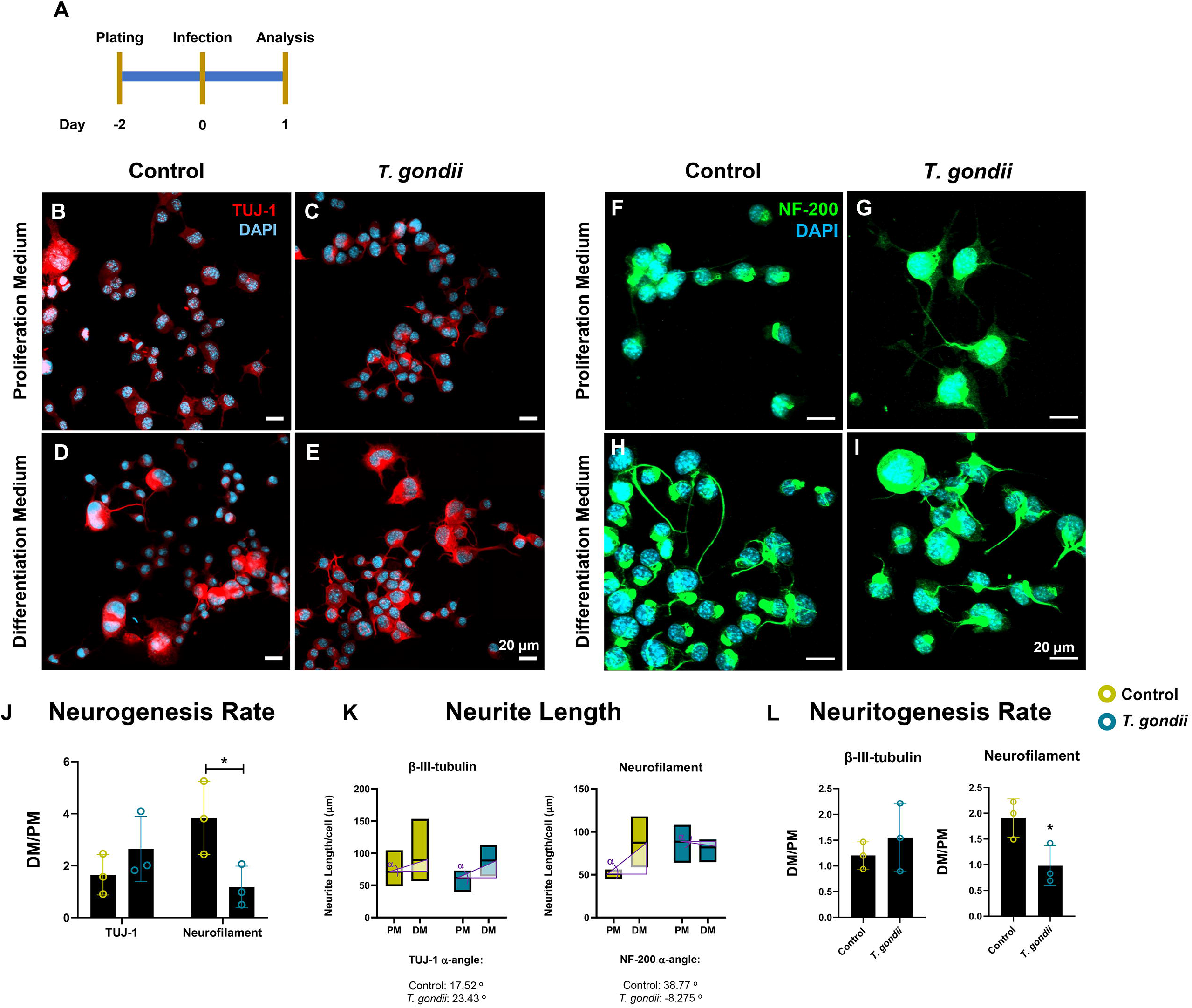
Neuritogenesis is affected by *T. gondii* infection in neuroblastoma cell line. Murine Neuro2a cells were infected in PM and half the cultures were switched to DM at 24 hpi (experimental design shown in **A**). Cells were stained with TUJ1 (**B**-**E,** in red) and NF-200 (**F**-**I**, in green) and neurogenesis rates were determined as the number of cells with neurites longer than the cell body in DM/PM (**J**). Host cell nuclei were stained with DAPI (in blue). Infection did not affect neurogenesis rates in TUJ1-stained cells but decreased significantly in NF-200-labeled cells. Neurite length was assessed in both TUJ1-or NF-200-positive cells (**K**). *T. gondii*-infected cultures showed no change in neuritogenesis rates in TUJ1, which were significantly decreased in NF-200-positive cells. *: p<0.05, unpaired Student’s T test of three independent experiments. Scale bars = 20 µm.

Next, we analyzed neurite outgrowth after 24 hours of induction with DM. Uninfected TUJ1 stained cells exhibited 71.25±29 µm long neurites in PM whereas cells in DM were 89.6±55 µm long, with an average neuritogenesis rate of 1.2±0.3 (α=17.52°). Proportionally to what was observed for neurogenesis rates, *T. gondii* infection did not affect neuritogenesis rates in TUJ1-positive Neuro2a cells, which displayed a DM/PM ratio of 1.55±0.7 (p=0.44, Unpaired T test). Regarding NF-200, uninfected Neuro2a cultures presented a 1.9±0.4 neuritogenesis rate, with PM-treated cells exhibiting 50.6±5 µm-long neurites, whereas cells treated with DM reached neurite length of 87.4±24 µm (α=38.77°). This effect was abrogated in *T. gondii*-infected cultures, which displayed a neuritogenesis rate of 0.97±0.4 (p=0.041, unpaired Student’s T test). Neurites in PM infected cultures were 88.5±22 µm long, whereas cells in DM exhibited 81.8±15 µm-long neurites (α= −8.28°, **Figure 4K-L**).

### 3. Astrogliogenesis is impaired in *T. gondii*-infected neurospheres

Astrocyte differentiation was evaluated by quantifying the number of glial fibrillary acidic protein (GFAP)-positive filaments in NPC cultures (**Figure 5**). At 48 h of plating, uninfected dishes contained 70.89±19 GFAP-positive filaments per mm, whereas *T. gondii*-infected exhibited 55.71±18 (p>0.05, **Figure 5B**, **C** and **F**). After 120 h of plating, the number of astrocytes increased in uninfected cultures, reaching an average of 102.2±34 GFAP filaments per mm (p=0.0048, Two-Way ANOVA with Bonferroni post-test, **Figure 5D**). However, 76.2±16.9 filaments/mm were detected in *T. gondii*-infected dishes (**Figure 5E**), which corresponded with a 25% reduction (p<0.05, Two-Way ANOVA with Bonferroni post-test). Insets **E’** and **E’’** highlight the intense parasitism of GFAP-positive cells as indicated by arrows in E’’, showing intracellular parasites.

**Figure 5:**
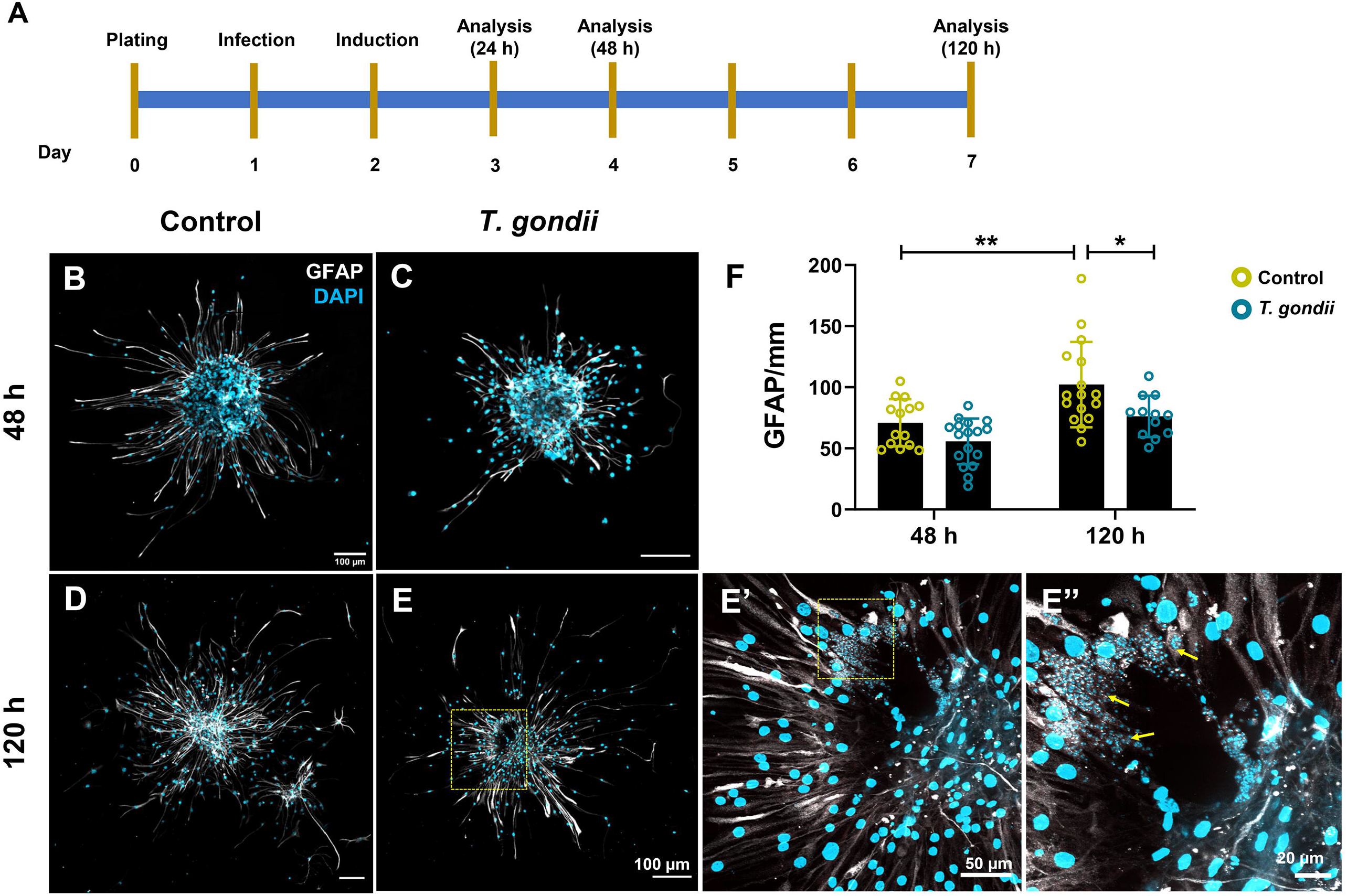
Astrocyte differentiation is impaired by *T. gondii*. Mouse NPCs were infected with *T. gondii* tachyzoites as floating neurospheres and then plated onto glass coverslips (the experimental design shown in **A**). Astrogliogenesis was assessed by GFAP (white) immunostaining after 48 (**B** and **C**) and 120 h (**D** and **E**) after plating. Cell nuclei are stained with DAPI (in blue) The number of GFAP-positive filaments was determined by confocal microscopy (**F**) and indicated that the number of GFAP-positive filaments increased over time (48-120 h). *T. gondii*-infected cultures exhibited decreased of GFAP-positive filaments at 120 h when compared to uninfected controls. Insets **E**’ and **E**’’ are higher magnifications of infected cultures at 120 h of plating (144 hpi), indicating intracellular parasites (arrows in E’’). *: p<0.05, **: p<0.01, Two-way ANOVA with Bonferroni post-test from at least 12 neurospheres obtained from a minimum of three independent experiments. Scale bars = 100 µm in **B**-**E**; 50 µm in **E’** and 20 µm in **E’’**.

### 4. Migratory potential is impaired in *T. gondii*-infected neurospheres

In order to assess the impact of *T. gondii* infection on the migratory ability of NPCs, cells were infected with tachyzoites one day after plating. At 24 hours after infection, neurospheres were plated onto glass coverslips coated with fibronectin and poly-L-lysine (**Figure 6A**). The area of the control neurospheres was of 488.3±372.5 µm^2^, whereas *T. gondii*-infected NF area was of 417.1±352.4 µm^2^, and no significant NF area changes were observed at this time (T = 0 h, **Figure 6B**) as determined by light microscopy. Cultures were then analyzed 24, 48 and 120 hours after plating (corresponding to 48, 72 and 144 hpi) and photographed by phase contrast microscopy (**Figure 6C**). The migration index was determined by the neurosphere areas (highlighted as yellow dashed lines in **Figure 6A**) and the perimeter was calculated using the ImageJ software. The cumulative migration index relative to the neurosphere area at T = 0 h was significantly reduced by *T. gondii* infection at 120 hours of plating (25.11±13.35 in controls versus 18.15±7.6 in infected cultures, p=0.032, Two-way ANOVA with Bonferroni post-test, **Figure 6C**). Interestingly, not only was the migrated area reduced in infected cultures, but we noticed that the radial migration pattern observed in uninfected neurospheres (as evidenced in black silhouettes in **Figure 6A**) was absent. Phase contrast microscopy images of adhered neurospheres suggest that infected cultures exhibit abnormal radial migration patterns in addition to reduced cell-occupied areas (**Figure 6A**). In order to investigate if migratory patterns were affected by infection, we performed Sholl analyses on nestin-, GFAP- and NF-200-stained cultures (**Figure 6D**). At 48 hpi, no changes were observed in migratory profiles, as indicated by the number of intersections among the radia (not shown). Nestin-stained neurospheres at 48 h exhibited decreased roundness and circularity and an increased aspect ratio (**Figure S2**). GFAP-labeled cultures displayed reduced roundness at 48 h, but no changes were detected in NF-200-stained cultures (**Figure S2**). After 120 h of plating, nestin-positive cells presented a reduced aspect ratio, whereas roundness and circularity (**Figure S2**), as well as the number of intersections per radius (**Figure 6E**), were not significantly altered. Regarding GFAP-positive cells, no changes were detected concerning roundness, circularity and aspect ratio at 120 h (**Figure S2**). However, *T. gondii*-infected cultures exhibited decreased intersections between 14 and 17 µm from the center of the neurosphere (p<0.001, Two-Way ANOVA with Bonferroni post-test, **Figure 6F**). *T. gondii* infection was more disruptive for NF-200-positive cell migration, since the number of intersections was diminished from the 13 to 36 µm-radia from the center of neurospheres (**Figure 6G**), although no changes were detected in roundness, circularity and aspect ratio (**Figure S2**).

**Figure 6:**
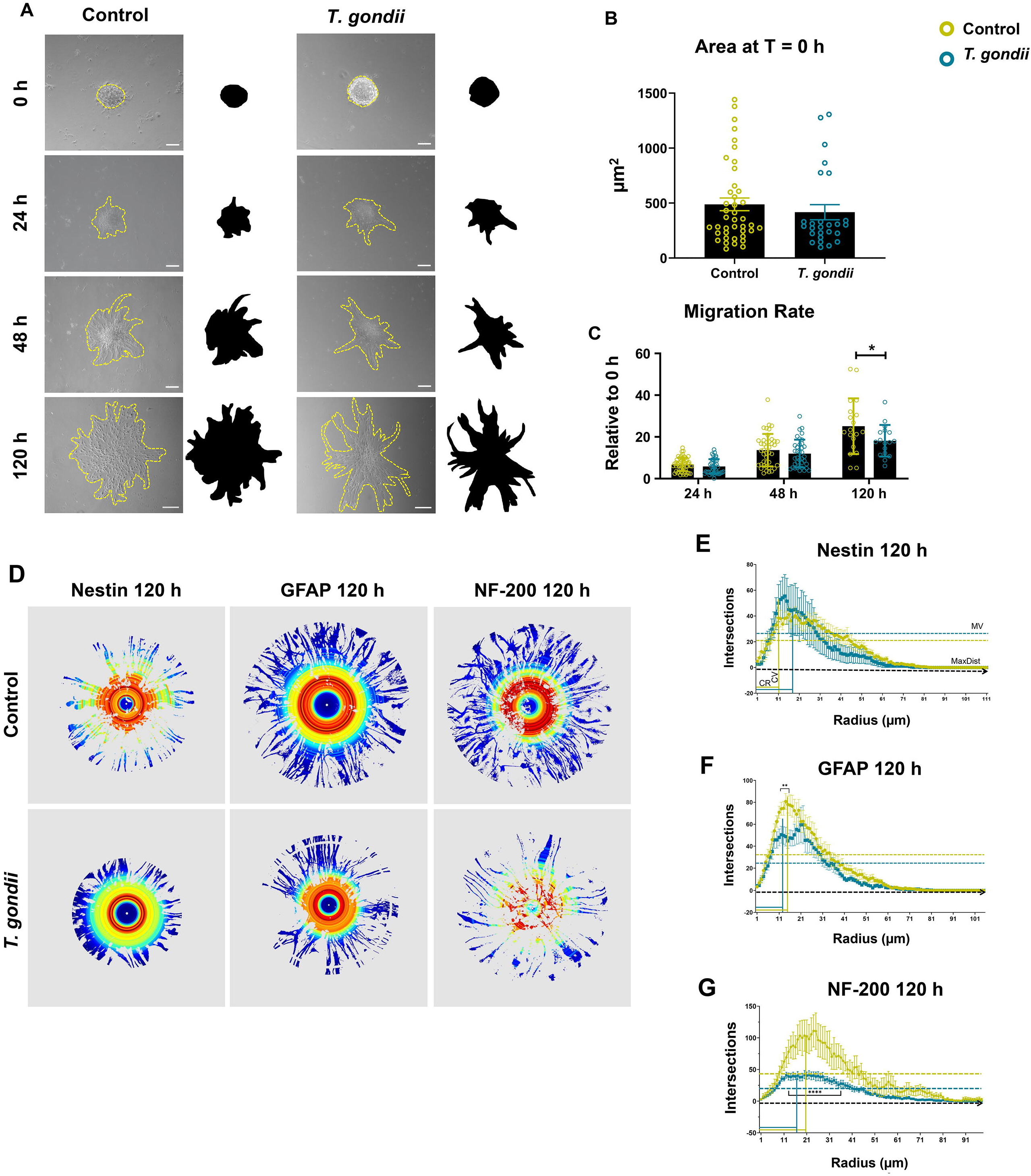
*T. gondii* infection impairs NPC migration patterns. Floating neurospheres were infected with *T. gondii* tachyzoites and adhered to glass coverslips, at 24 hpi. Cultures were imaged by phase contrast light microscopy at different times to assess migration (**A**). At 0 h, no changes were seen in neurosphere sizes (**B**). Migration rates were determined by cell-occupied perimeter at 24, 48 and 120 hours in relation to the correspondent area at 0 h (**C**). *T. gondii* infection significantly impaired migration at 120 h. Radial migration patterns were evaluated by Sholl plugin in cultures stained for different neural populations at 120 h (**D**). Nestin-positive cells (**E**) showed no significant migration alterations, whereas GFAP-(**F**) and NF-200-stained cells (**G**) exhibited altered parameters. **CM**: Critical Value, **CR**: Critical Radius; **MV**: Mean Value. Scale bars: 10 µm for 0 h; 20 µm for 24 and 48 h; 100 µm for 120 h.

## DISCUSSION

The CNS becomes established upon the formation of the neural tube (O’Rahilly & Muller, 2010; Devakumar et al., 2018). The neural tube contains an inner surface, named the ventricular zone, located adjacent to the ventricle. The outer surface is covered with a basal membrane and the pia mater, thus named the pial surface. Cell divisions occur on the ventricular or apical surface of the neuroepithelium (Reviewed by Garcia-Moreno & Molnar, 2020). The embryonic brain is initially composed entirely of proliferative neuronal progenitors, which reside within the ventricular zone that borders the neural tube lumen. However, with subsequent development, neurons begin to emerge and a new population of deeper subventricular zone (SVZ) neural progenitors arises. Proliferation of SVZ cells contributes to further expansion of the brain’s neuronal population. These processes are finely orchestrated by cellular and molecular events, which can lead to cortical malformations when disturbed by teratogenic agents.

*T. gondii* is part of the TORCH group of microcephaly-causing pathogens when transmitted during pregnancy. Cellular and molecular effects of *T. gondii* infection during pregnancy in humans are, up to now, still poorly understood phenomena, given the difficulties in performing clinical studies in pregnant women. In the particular case of the Brazilian public health system, intrinsic complications are associated to accurate diagnosis and treatment accessibility. Such difficulties indicate an urgent need for the development of experimental models that mimic neural development to allow the investigation of cellular and molecular events triggered by *T. gondii* infection on NPC biology, both *in vitro* and *in vivo*.

In the present study, we used a primary murine NPC culture exhibiting neuronal and glial multipotency. Cells were maintained in a PM containing recombinant EGF, which maintained the cells in a proliferative, non-adherent stage (Santiago et al., 2010). We assessed their *in vitro* proliferation, migration and potential to differentiate into astrocytes and neurons upon removal of EGF and addition of pro-neurogenic factors (N2 and retinoic acid). Infected neurospheres displayed reduced proliferation at 48 and 72 hours post infection, which resulted in diminished cellularity at 96 hpi. Reduced proliferation may decrease the pool of neural stem cells, that must proliferate and eventually perform asymmetric or symmetric divisions to generate neurons or renew NSC pools, respectively (Uzquiano et al., 2018). The fact that significant decreases in Ki67-positive cells were restricted to the two intermediate time points may reflect active tachyzoite proliferation, which may be attenuated after tachyzoites convert spontaneously to bradyzoites. *In vitro* stage conversion has been shown to occur in neural cell cultures (Lüder et al., 1999) and in our neurosphere cultures, bradyzoite-specific BAG-1 mRNA have been detected at 96 hpi by RT-PCR (Adesse, D., personal observation). Our group recently described that radial glial cell infection with *T. gondii* tachyzoites (Me49 strain, the same utilized herein), led to proliferation reduction with no apoptosis induction (Marcos et al., 2020) of these early neural stem cells. Therefore, reduced proliferation in infected dishes could contribute to reduced neurogenesis in infected animals. Interestingly, it has been recently described that *T. gondii* disrupts correct cytokinesis patterns and the formation of the mitotic spindle in bovine endothelial cells (Velásquez et al., 2019). It is known that changes in spindle structure, with or without cell cycle protein alterations, can lead to abnormal retinal and brain cortex development (Kitawaga et al., 2011; Uzquiano et al., 2018), which could also explain how CT affects these processes.

We then plated both infected and uninfected neurospheres on a substrate containing fibronectin and observed neural population migratory profiles and differentiation. The reduced migration rate of nestin-positive cells observed in infected cultures may be, in part, caused by changes in Extracellular Matrix (ECM) components or focal adhesion signaling pathways. It has been reported that *T. gondii*-infected astrocytes cultures exhibit matrix metalloproteinase-2 and −9 overexpressions through ERK/NF-κB pathways (Lu & Lai, 2013a), which, in turn, degrade fibronectin (Lu & Lai, 2013b). Moreover, *T. gondii* possesses an aminopeptidase N that is a member of the M1 family of metalloproteases (Li et al., 2017) and which could also play a role in ECM remodeling. Accordingly, ECM components can also be involved in neuritogenesis, as recently reported by Sugahara and colleagues (2019) in Neuro2a cells, in which vitronectin and β5 integrin were shown to be involved in cell polarization and neurite outgrowth. Neuroblast migration is an important step during *in vivo* cortical neurogenesis in mice and migration defects can contribute to cortical malformations, including microcephaly (Garcez et al., 2015; Nicole et al., 2018). *T. gondii* infection also displays the ability to recruit microtubules to the vicinity of its parasitophorous vacuole (Andrade et al., 2001; Cardoso et al., 2016; Paredes-Santos et al., 2018). Cytoskeleton proteins such as tubulin play a major role in neuroblast migration and cortical neurogenesis (Belvindrah et al., 2017; reviewed by Francis & Belvindrah, 2018), thus indicating that infected progenitor cells may indeed display migratory defects.

After two or five days of migration on a fibronectin substrate and with a culture medium without EGF and supplemented with N2 and retinoic acid, cells began to differentiate into astrocytes and neurons, as detected by GFAP and neurofilament expressions. We observed that infection led to no changes in proliferation rates and proportion of progenitor cells, as revealed by Ki67 and nestin staining, respectively. However, the number of progenitors (Ki67+/nestin+) was reduced in uninfected cultures at 120 h when compared to 48 h as a natural consequence of cell differentiation, thus decreasing this population. In our model, no evident cell death was detected, as revealed by pyknotic nuclei counts by DAPI staining. This contrasts with what was previously reported when C17.2 NSCs were infected with the *T. gondii* RH strain (Wang et al., 2014). Parasite backgrounds seems to play a crucial role in their interaction with neural stem cells, since a Chinese *T. gondii* isolate with a peculiar genotype. exhibiting type I and II background features induced weaker cell death in these cultures (Zhou et al., 2015). Indeed, the expression rates of *T. gondii* effector proteins, such as ROP16 and ROP18, vary among genetic groups I, II, III and can display multiple combinations in atypical strains of wild isolates (Shwab et al., 2014; Shwab et al., 2016), which confer different degrees of virulence. However, in our study we used Me49, a reference strain of the Type II parasite, which does not express high levels of ROP16 (Saeij et al., 2007; 2006), and could, in part, explain the low rates of host cell lysis and exclude possible regulation by these two main effector proteins.

Regarding neuronal production, a trend of increasing TUJ1-positive cells at 48 h was observed in attached neurospheres (as well as in Neuro2a cells), although no statistical significance was detected. As neurogenesis progression occurred from 48 to 120 hours, the number of TUJ1+ cells drastically decreased in both uninfected and infected dishes. Neuronal maturation progression was observed by NF-200 staining, where control cultures displayed an increase in this marker from 48 to 120 h, thus confirming neurogenesis. This effect was inhibited in *T. gondii* infected cultures. Impaired neurogenic potential (as estimated by TUJ1+ cells) was previously demonstrated by our group in radial glia cell cultures infected with Me49 tachyzoites (Marcos et al., 2020). This effect was also observed when C17.2 neural stem cell line cells were treated with soluble factors released from *T. gondii* (Gan et al., 2016), in a process mediated by the Wnt/β-catenin signaling pathway. Our group has also described that direct infection of skeletal muscle precursors (myoblasts) affects Wnt/β-catenin signaling pathway activation, which impairs myogenesis and leaves cells in a proliferative, undifferentiated state (Vieira et al., 2019).

In order to further understand the effect of *T. gondii* infection on neuronal maturation, we used Neuro2a cells. Upon serum withdrawal, these cells activate EGFR, ERK and Akt signaling pathways that promote *in vitro* neuronal differentiation (Evangelopoulos et al., 2005). When we infected Neuro2a neuroblasts with *T. gondii* tachyzoites in PM and then exchanged them to DM, a significant reduction in neuron-like cells, with reduced neurites was observed when compared with control cultures. In addition, the same trend of increasing TUJ1+ cells as seen in neurospheres was observed, suggesting that infection may lead to interruption (or, at least, a delay) in TUJ1 to NF-200 progression. Neurite outgrowth has been evaluated in Neuro2a cells by other groups under different stimuli/treatments, and it has been reported that this process can be modulated by carbazole derivatives (Furukawa et al., 2019), miRNA (You et al., 2020), Mob proteins (Lin et al., 2011) and polyphenols, including Resveratrol and Apigenin (Namsi et al, 2018). Interestingly, Resveratrol has been shown to revert *T. gondii*-induced cytokine release and purinergic signaling imbalance in neural progenitor cells (Bottari et al., 2019). Other groups have previously utilized neuroblastoma cell lines to investigate the impact of *T. gondii* infection, where transfection of human neuroblastoma SH-SY5Y cells with ROP16 effector protein led to apoptosis and cell cycle arrest (Chang et al., 2015), which also resulted in the remodeling of host cell transcriptomic networks, including those related to nervous system development, apoptosis and transcriptional regulation (Fan et al., 2016). Our results demonstrating reduced neuritogenesis are also in accordance to what was recently shown in infected cultures of superior cervical ganglion cells, in which neurite networks were decreased in cultures infected with a highly virulent strain of *T. gondii* (TgCTBr9, genotype #11) from 48 to 192 hpi (Barbosa et al., 2020).

Another interesting *T. gondii* feature is the fact that this parasite possesses a tyrosine hydroxylase ortholog, the enzyme responsible for producing dopamine, thus increasing neural cell dopamine production and metabolism (Prandovzky et al., 2011; Martin et al., 2015). Neurogenesis can rely on dopaminergic signaling and reduced neurogenesis has been detected in autopsied brains from Parkinson disease patients and dopaminergic dysfunction animal models (Hoglinger et al., 2004; Berg et al., 2013). As precursor cells in the SVZ, including neuroblasts, express dopamine receptors, it is conceivable that dopamine may control neurogenesis aspects in this region and that *T. gondii* infection can imbalance this finely tuned process (Diaz et al., 1997; Lopatina et al., 2019). Since parasitism of host cells was restricted to a few cells within a single neurosphere in our infection model, it is reasonable to assume that soluble factors secreted by infected cells may also play a paracrine role in the deleterious effects described herein.

Finally, astrogliogenesis, as revealed by GFAP immunostaining was significantly reduced in infected cultures. This contrasts with what we recently described in radial glia culture infection, in which GFAP+ cells remained unaltered (Marcos et al., 2020). Contrasting results regarding neuronal and astrocytic differentiation in radial glia and intermediate progenitors due to *T. gondii* infection may, in fact, reinforce the differences in clinical outcomes when vertical transmission occurs during different gestational periods. Moreover, our group has shown, using two independent systems, that *T. gondii* decreases the secretion of transforming growth factor beta 1 (TGF-β1), in both radial glia (Marcos et al., 2020) and skeletal muscle cells (Vieira et al., 2019). TGF-β1 is a pleiotropic cytokine involved in organogenesis and pathogenic events in vertebrates (reviewed by Massagué, 1998). Regarding cortical development, TGF-β1 is crucial for astrogliogenesis in mice, and treatment with TGF-β1 inhibitor SB431542 results in reduced astrocyte populations in the cerebral cortex (Stipursky et al., 2014).

Taken together, our data point to a deleterious *in vitro T. gondii* effect on the biology of neural progenitor cells, including proliferation, migration patterns and neurogenic and gliogenic potential. This model will serve as basis for additional mechanistic studies concerning these phenomena and for the elucidation of cellular events that may take place in the developing mouse brain during congenital infection.

## ACKNOWLEDGMENTS

The authors thank Mrs. Sandra Maria Oliveira de Souza (LBE, IOC) for excellent technical support; Mrs. Thalyta Priswa Andrade for help in generating initial neurosphere assays; Mrs. Camilla Bayer (IBCCF-UFRJ) for help with confocal imaging and Prof. Joice Stipursky (ICB-UFRJ) for critical reading and result discussions. This work was supported by Fundação Oswaldo Cruz (Edital INOVA Geração de Conhecimento 2018, grant number 3231984391), Conselho Nacional de Pesquisa e Desenvolvimento Tecnológico (CNPq, grant numbers: 401772/2015-2 and 444478/2014-0 for DA), Fundação Carlos Chagas Filho de Amparo à Pesquisa do Rio de Janeiro (FAPERJ, Projetos Temáticos grant number SEI-26 260003/001351/2020 and Redes em Saúde, grant number E-26-211.570/2019 for D.A and H.S.B.). L.B.P. is sponsored by a Ph.D. fellowship from CAPES/Brazil.

## AUTHOR’S CONTRIBUTION STATEMENT

**LBP:** performed the experiments, carried out the immunocytochemistry assessments, confocal analyses, morphometrical analyses and statistical analyses and wrote the manuscript draft; **HSB:** contributed with reagents, equipment and critical discussions regarding the experimental design and assisted with the Electron Microcopy analyses; **MFS:** established the mouse neurosphere model and assisted in the study conceptualization and data interpretation; **DA:** was involved with the project’s conceptualization and coordination, performed the statistical analyses and prepared the manuscript and figures.

---

**Supplementary Figure S1: *T. gondii* infection does not increase pyknosis in mouse NPCs.** Migrated neurospheres were fixed at 48 h after plating, stained with SAG-1 antibody (in red) and counterstained with DAPI (in blue). Pyknotic nuclei are indicated by arrows and insets exhibit higher magnifications of corresponding images. No significant pyknosis changes were detected in *T. gondii*-infected cultures compared to uninfected dishes. Scale bars = 20 µm.

**Supplementary Figure S2: Sholl analysis of nestin-, GFAP- and NF-200-stained cultures.** Additional parameters such as roundness, circularity and aspect ratio were determined using Sholl plugin of plated neurospheres after 48 and 120 h. *: p<0.05; **: p<0.01.

