## Supplementary figures and images for "Infection of mouse neural progenitor cells by *Toxoplasma gondii* affects *in vitro* proliferation, differentiation and migration"

### Supplementary Figure 1

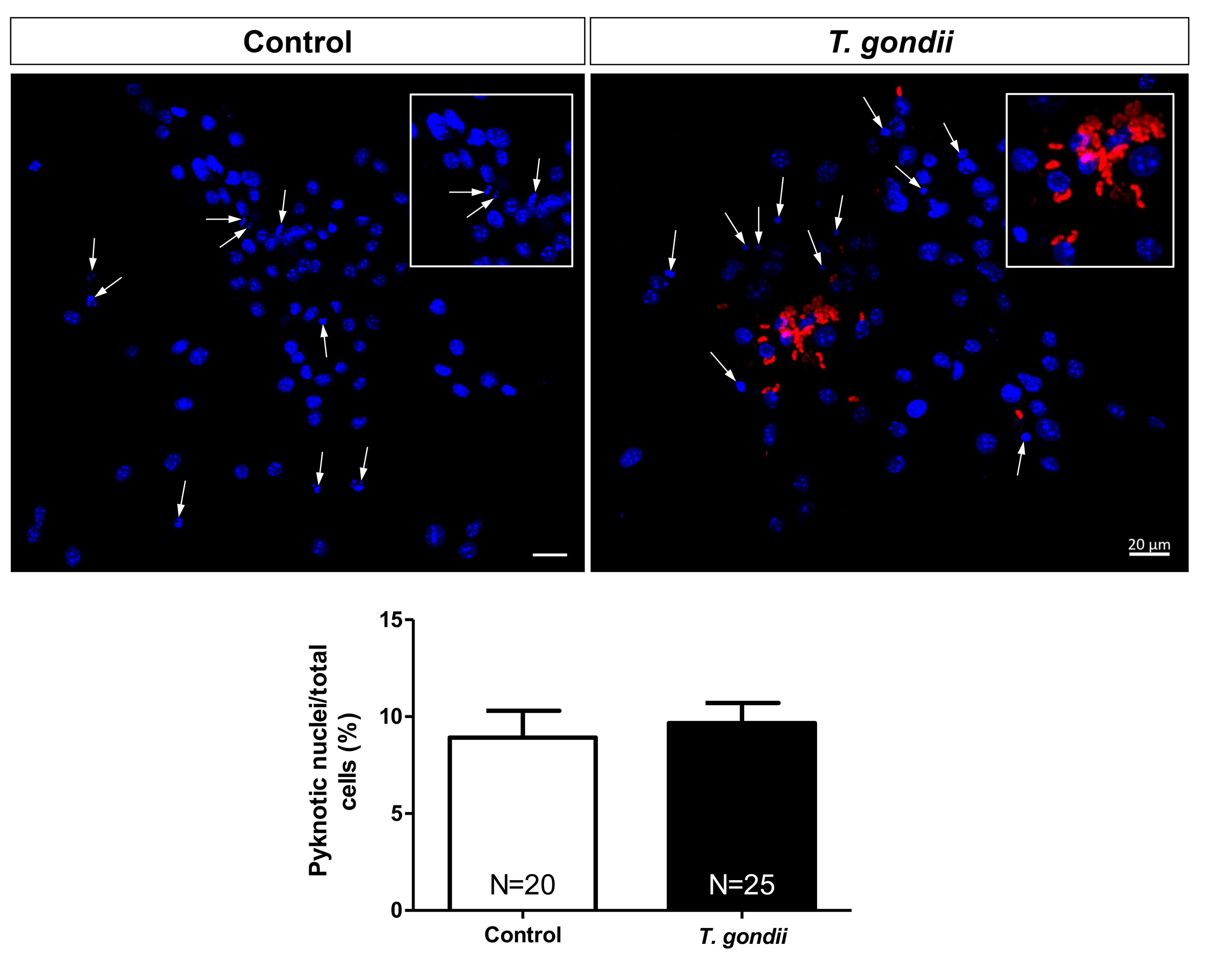

### Supplementary Figure 2

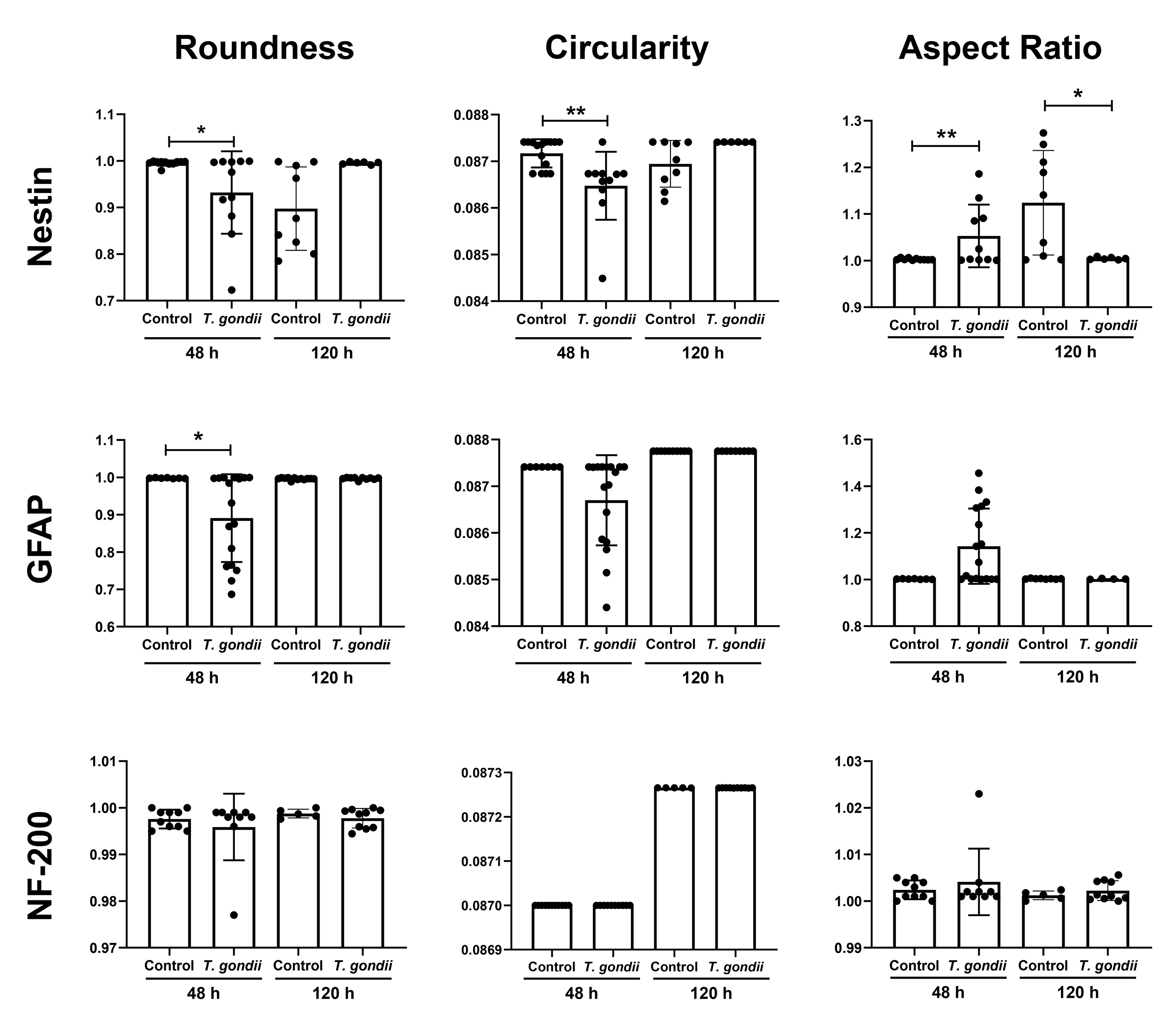
